# Obesity and sarcopenia affect the metabolite profiles of pet dogs

**DOI:** 10.1101/2025.03.31.645884

**Authors:** Claudia Elo, Mirja Kaimio, Essi Leminen, Hannes Lohi

## Abstract

Overweight and reduced muscle mass significantly affect canine health, yet their metabolic implications in dogs require further investigation. This study aimed to characterize metabolic alterations associated with overweight, obesity, and reduced muscle mass in dogs. An observational, cross-sectional study was conducted to evaluate differences in metabolite concentrations among pet dogs with varying body and muscle conditions. Dogs older than two years were included in the Body Condition Score (BCS) analysis (n=186), while only dogs older than eight years were included in the Muscle Condition Score (MCS) analysis due to age-related sarcopenia (n=99). Metabolomic analyses were performed using a validated, canine-specific nuclear magnetic resonance (NMR) platform. Associations between metabolite concentrations and body or muscle condition were examined independently using generalized linear models, adjusting statistically for age and health status. Increasing overweight status correlated with notable disruptions in lipid and glucose metabolism, alongside elevated inflammation markers. Specifically, several lipid metabolites increased linearly with increasing overweight, as did citrate, lactate, pyruvate, and the inflammatory biomarker GlycA. Conversely, decreased muscle mass showed relatively modest independent metabolic effects, including elevated GlycA, an increased phenylalanine-to-tyrosine ratio, and alterations in VLDL subclass composition. Muscle mass was closely associated with age and overall health status, with health status emerging as the primary determinant of metabolite concentrations. Moderate to severe muscle loss was predominantly observed in dogs with underlying diseases. The metabolic consequences of overweight and sarcopenia, particularly increased inflammation common to both conditions, could significantly impact a dog’s overall health. These findings reinforce existing evidence regarding the detrimental health effects associated with overweight and sarcopenia in dogs. Furthermore, the study suggests that underlying disease should be suspected in older dogs presenting highly abnormal metabolite profiles or moderate to severe muscle loss.

## INTRODUCTION

Body condition and muscle mass significantly determine and reflect health and wellbeing in pet dogs. Similar to trends observed in humans, the prevalence of overweight and obesity among dogs has notably increased in recent years, becoming a critical health concern ^1^. Concurrently, muscle loss, particularly in older or diseased dogs, is common and adversely affects wellbeing, daily performance, and lifespan ^2^. Nevertheless, obesity and muscle loss remain considerably underestimated issues, despite their substantial negative impact on affected animals.

Obesity is a complex condition characterized by body weight exceeding optimal weight by at least 15%, primarily due to excessive accumulation of adipose tissue ^3^. Estimates indicate that the prevalence of canine overweight and obesity ranges from 10% to 59% across the general dog population ^1,3–6^. Obesity typically arises from energy imbalances resulting from high caloric intake combined with insufficient physical activity. Various contributing factors include specific diseases, medications, neutering, feeding habits, activity levels, owner influences, and genetic predispositions ^1,7–11^. Obesity is linked to multiple health complications, such as cardiovascular, respiratory, endocrine, orthopedic, and oncological disorders ^12–14^, as well as reduced lifespan ^15–17^. Additionally, obesity negatively impacts canine welfare and quality of life ^18^. Given the high prevalence and welfare implications of obesity, a comprehensive understanding of its metabolic effects is essential for improved prevention and management strategies.

Muscle loss and its negative impacts on human health have long been acknowledged; however, veterinary medicine has only recently begun to routinely assess cachexia and sarcopenia. Cachexia refers to muscle wasting resulting from chronic diseases such as chronic kidney disease (CKD), congestive heart failure (CHF), or cancer, characterized by significant loss of lean body mass ^2^. Sarcopenia, in contrast, is an age-related condition involving the loss of lean body mass independent of underlying disease. Due to potential increases in body fat mass, sarcopenia may not be easily noticeable visually, making it difficult to detect.

In humans, sarcopenia has been associated with various physiological factors, including decreased growth hormone and testosterone levels, reduced protein synthesis, insulin resistance, increased cytokine production, and physical inactivity ^19,20^. Both cachexia and sarcopenia negatively affect muscle strength, immune function, quality of life, and mortality in humans. While research in pets remains limited, a few feline studies indicate that weight loss is associated with increased mortality ^21–23^. Additionally, weight loss and weakness in ill or elderly animals often significantly influence euthanasia decisions ^24^. Despite extensive research on the metabolic consequences of human sarcopenia, similar studies have not yet been conducted in dogs. Therefore, understanding the molecular and metabolic changes during canine sarcopenia is essential.

A promising approach to detect systematic metabolic disruptions and potential biomarkers associated with obesity and muscle loss in dogs is a recently developed and clinically validated nuclear magnetic resonance (NMR) spectroscopy-based metabolomics platform ^25^. This platform measures 123 metabolites, including creatinine, albumin, glucose, glycolysis-related metabolites, GlycA, amino acids, cholesterol, triglycerides, fatty acids, and provides comprehensive lipoprotein analysis—all derived from a single plasma or serum sample in one measurement. Many of these metabolites have been previously identified or hypothesized to be affected by obesity and/or muscle loss in dogs or other species.

This study aims to characterize the metabolic alterations associated with i) overweight/obesity and ii) sarcopenia in dogs. Understanding the metabolic consequences of suboptimal body composition and muscle condition contributes significantly to our knowledge of their detrimental health effects. We anticipate identifying clinically relevant metabolic changes that can substantially enhance the monitoring, achievement, and maintenance of optimal body weight and muscle mass in dogs.

## MATERIALS AND METHODS

We conducted an observational, cross-sectional study to investigate differences in metabolite concentrations among dogs with varying body condition scores (BCS) and muscle condition scores (MCS). Dogs were recruited between May 2022 and July 2023 from two veterinary hospitals in Finland: Animal Hospital Evidensia Tammisto in Vantaa and Espoo Veterinary Hospital in Espoo. Recruitment primarily occurred during regular patient visits or through study advertisements. Our goal was to enroll at least 20 dogs in each BCS and MCS category.

Female dogs that were in heat, pregnant, lactating, or pseudo pregnant were excluded due to known metabolic effects associated with these conditions ^26,27^. Additionally, dogs belonging to sighthound breeds were excluded because they exhibit significant metabolic differences compared to other breeds ^28^.

Initially, we planned to enroll dogs aged between 10 and 14 years for the MCS analysis, requiring them to have no internal diseases based on history and physical examinations within the previous three months. However, it quickly became apparent that meeting these criteria was challenging due to the prevalence of diseases among geriatric dogs and the reluctance of owners to bring healthy geriatric dogs to veterinary visits. Consequently, we expanded our inclusion criteria to dogs aged eight years or older, provided they were free from diseases known to significantly impact metabolite concentrations, such as endocrine disorders, liver or renal failure, or other severe illnesses.

For the BCS analysis, we initially targeted dogs between two and six years of age that were deemed healthy based on recent veterinary assessments. However, overweight and obesity were uncommon in younger adult dogs within this population, whereas they were frequently observed among senior dogs. Therefore, the age criteria were adjusted to include all dogs aged two years or older, applying the same health criteria as for the MCS analysis.

All enrolled dogs were weighed, and their BCS was evaluated using a 1–9 scale. MCS was assessed visually and through palpation, based on the World Small Animal Veterinary Association (WSAVA) guidelines ^29,30^, which classify muscle condition from normal to severe muscle loss. As the original WSAVA guidelines provide only illustrative drawings without written descriptions, we developed written descriptions for each MCS category specifically for this study. These descriptions were reviewed and approved by a member of the WSAVA Global Nutrition Committee.

BCS and MCS assessments were performed by two qualified veterinary nurses who received specialized training prior to conducting evaluations. Additionally, the nurses conducted comprehensive interviews with dog owners to gather detailed demographic, health, and dietary information about their pets.

Blood samples for nuclear magnetic resonance (NMR) analysis were collected via cephalic venipuncture, drawing 2 ml of blood into Vacuette LH Lithium Heparin PREMIUM tubes (2 ml capacity). Samples were immediately centrifuged, and aliquots ranging from 110 to 500 µl of heparinized plasma were separated. These plasma samples were promptly cooled and refrigerated for up to one week before transportation on ice packs to PetMeta Labs in Helsinki, Finland, for NMR analysis. Upon arrival, samples were stored at −80°C and subsequently analyzed in batches of 94. Most dogs participating in this study were concurrently enrolled as healthy controls in a related study evaluating metabolic changes associated with inflammatory diseases. Clinical chemistry, hematology, and C-reactive protein (CRP) analyses were conducted for these dogs. A veterinarian thoroughly reviewed all clinical and laboratory data to confirm the health status of each dog. Thirteen dogs were excluded based on abnormal findings from clinical chemistry, hematology, or CRP tests. Specific reasons for exclusion included elevated liver enzymes indicative of liver disease, anemia or inflammatory cell alterations, and elevated CRP levels. Additionally, one dog was removed due to a health history of canine epileptoid cramping syndrome, chronic itching, tracheal hypoplasia, and thrombocytopenia.

The final cohort of 285 dogs consisted of 186 dogs in the BCS study and 99 dogs in the MCS study. Clinical chemistry, hematology, and CRP data were available for 172 (92%) of the dogs in the BCS group. Among the 186 BCS dogs, 37 had identified health conditions, while 29 out of the 99 MCS dogs had similarly diagnosed conditions. The most frequently diagnosed conditions in both groups were osteoarthritis, spondylosis, and periodontitis.

### ^1^H NMR spectroscopy

The metabolomics approach employed in this study is a clinically validated, targeted ^1^H nuclear magnetic resonance (NMR) spectroscopy method specifically optimized for canine samples ^25^. A similar methodology has been extensively applied in human studies, with detailed descriptions of this highly automated process available in previous publications ^25,31,32^. Briefly, the method utilizes a Bruker AVANCE III HD 500 NMR spectrometer equipped with a 5 mm triple-channel (1H, 13C, 15N) z-gradient Prodigy probe head, a SampleJet automated sample changer (Bruker Corp., Billerica, Massachusetts, USA), and a PerkinElmer JANUS Automated Workstation featuring an 8-tip Varispan dispensing arm (PerkinElmer Inc., Waltham, Massachusetts, USA) ^31^.

Sample preparation for this method is straightforward, requiring minimal handling. It involves gentle mixing of samples, removal of any precipitates by centrifugation, transferring each sample into individual NMR tubes, and subsequent mixing with sodium phosphate buffer ^31,33^.

NMR spectra processing is conducted using scripts specifically optimized for canine samples, with metabolite quantification provided in absolute units through regression modeling ^25,32^. Integrated quality control procedures ensure the reliability of the analysis via proprietary software ^32^. The targeted approach quantifies 123 metabolites, including creatinine, albumin, glucose, various glycolysis-related metabolites, GlycA, amino acids, cholesterol subtypes, triglycerides, fatty acids, and provides comprehensive lipoprotein profiling ^25^.

### Statistical analyses

Data processing and statistical analyses were performed using the R programming environment ^34^ and Microsoft Excel (Microsoft Office, Redmond, WA, USA). One dog with a BCS of 3 was excluded from the BCS analysis. Due to limited sample sizes, dogs with a BCS of 9 (n=4) were combined with those with a BCS of 8, forming a group representing severe obesity (BCS ≥8). Similarly, dogs with severe muscle loss (MCS4, n=2) were merged with those exhibiting moderate muscle loss (MCS3). Initially, metabolite data were reviewed for data integrity and missing values. Four dogs were excluded due to predominantly missing metabolite results, attributable to solid material in the tubes (n=3) and a technical error (n=1). Metabolites with more than 20% missing values, specifically XL-VLDL-PL, XL-VLDL-CE, and XL-VLDL-FC, were excluded from further analysis. Remaining missing values were imputed using random forest imputation via the missForest version 1.5 ^35^, dplyr version 1.1.2 ^36^, and tidyverse ^37^ packages. The out-of-bag (OOB) error of 0.001 indicated excellent imputation performance. Metabolite data were also screened for outliers, but no additional exclusions were necessary.

Due to high intercorrelation among metabolite variables, the significance threshold was adjusted using principal component analysis (PCA). PCA was conducted using FactoMineR ^38^ and factoextra version 1.0.7 ^39^ packages. Five principal components accounted for 95% of the data variance, leading to a Bonferroni-corrected significance threshold of 0.01 (0.05/5).

We assessed potential confounding variables by examining differences in age, sex, sterilization/castration status, fasting duration, BCS/MCS, and health status between groups. Age differences were analyzed with the Kruskal-Wallis test and Dunn’s post-hoc test (FSA package ^40^), while Chi-squared tests were applied for other physiological characteristics (stats package ^34^). The significance threshold was set at 0.05. Age significantly differed between MCS groups, while health status differed between both BCS and MCS groups; thus, age and health status were accounted for in subsequent analyses. Dietary and exercise habit differences between groups were evaluated with Chi-squared tests to explore environmental factors influencing BCS and MCS. Heatmaps illustrating metabolite concentrations by MCS and BCS were generated using ComplexHeatmap ^41^, circlize ^42^, tidyverse ^37^, and viridis version 0.6.4 ^43^ packages.

Generalized linear models (GLMs), adjusted for age and health status, were utilized to evaluate the independent effects of BCS/MCS on metabolite concentrations. GLM assumptions—including dispersion, linearity of age, outliers, residuals, autocorrelation, and multicollinearity—were carefully validated. For metabolites demonstrating nonlinear age effects, quadratic age terms were included. Packages employed for model construction and validation included ggplot2 ^44^, devtools version 2.4.5^45^, patchwork version 1.1.2 ^46^, gplots version 3.1.3 ^47^, dplyr version 1.1.2 ^48^, boot version 1.3-28.1 ^49^, rcompanion version 2.4.30 ^50^, ggpubr version 0.6.0 ^51^, gam version 1.22-2 ^52^, splines ^34^, foreach version 1.5.2 ^53^, broom version 1.0.4 ^54^, car ^55^, and stats ^34^. All model assumptions were met for the BCS analysis. Individual contributions of BCS, age, and health status to metabolite concentrations were visualized using effect plots from the effects package ^55^.

Due to significant multicollinearity among age, health status, and MCS in the MCS analysis, models with only age, only health status, only MCS, and combinations thereof were compared using the Akaike information criterion (AIC). The model with the lowest AIC was selected as the best explanatory model for metabolite differences. The resulting p-values were visualized via heatmaps using the ComplexHeatmap package ^41^.

### Ethical approval

Blood sampling from privately owned pet dogs was approved in the project license by the Animal Ethical Committee of the County Administrative Board for Southern Finland (ESAVI/16933/2021). Whenever possible, blood samples for this study were obtained simultaneously with routine laboratory diagnostic sampling conducted for the dog’s clinical benefit, thus minimizing additional stress. All dog owners provided informed consent before participating in the study and were free to withdraw at any time. Dog information was recorded using pseudonymization to maintain confidentiality, and owner information was neither collected nor used in the study.

## RESULTS

### Body condition score Demography

The demographical information of dogs included in the BCS analysis is presented in Table 1. The median age was on the senior side in all BCS groups, although ages ranged from young adult dogs to geriatric ones in all groups. All groups had slightly more females than males. Health status was the only demographical parameter exhibiting significant differences between the groups. All groups consisted of various breeds, but sighthounds were excluded per study design.

**Table 1.** Demographical information of the 186 dogs included in the BCS study. * significant (Chi-square p < 0.05) difference between groups.

| BCS | N | Age median (range) | %M/F | % castr | % ster | Fasting% <12h/ >12 h | % disease* | N breeds |
| --- | --- | --- | --- | --- | --- | --- | --- | --- |
| 4 | 18 | 9.9 (3.1 – 14.7) | 44/56 | 25 | 80 | 50/50 | 5.6 | 16 |
| 5 | 103 | 8.3 (1.9 – 16.3) | 39/61 | 22 | 44 | 46/54 | 17.5 | 51 |
| 6 | 32 | 7.9 (2.1 – 16.4) | 41/59 | 62 | 47 | 44/56 | 15.6 | 19 |
| 7 | 18 | 7.5 (2.7 – 13.7) | 50/50 | 67 | 56 | 22/78 | 33.3 | 16 |
| 8-9 | 15 | 8.0 (2.2 – 12.4) | 47/53 | 71 | 50 | 53/47 | 33.3 | 12 |

No significant differences (chi-squared test, p < 0.05) were found regarding owner-reported daily exercise duration (hours of outdoor activity), exercise type (on-leash, off-leash, or both), food type (commercial diet, raw diet, combined commercial and raw diet, home-cooked diet, or other), daily feeding frequency (once, twice, three or more times, ad libitum, or other), treat frequency (daily, several times per week, monthly, yearly, or never), or oil supplementation frequency (daily, several times per week, monthly, yearly, or never). However, a significant difference was observed concerning how daily food portions were determined (food package instructions, veterinarian’s recommendations, visual/palpational observation of BCS by the owner, based on the dog’s hunger status, ad libitum feeding, other). Specifically, obese dogs were unique in having their daily food amounts set according to veterinarians’ recommendations. In contrast, most other dogs had their food portions determined by the owner’s visual or palpational assessment of body condition.

### Effects of body condition on metabolism

Overweight and obesity resulted in linear alterations in lipid metabolism, glucose metabolism, and levels of the inflammatory marker GlycA (Figure 1, Supplementary Table). Among the glucose-related metabolites studied, citrate, lactate, and pyruvate concentrations significantly increased with rising body condition scores (BCS); however, glucose levels did not show statistically significant changes. Lipid metabolism changes primarily involved alterations in HDL, VLDL, total fatty acids, and triglycerides. Specifically, with higher BCS, HDL particles decreased in size, became more numerous (especially small HDL particles), and exhibited elevated triglyceride content. Furthermore, overweight dogs displayed disruptions in triglyceride metabolism characterized by elevated triglyceride levels, increased VLDL concentrations, and larger VLDL particles, particularly within the larger VLDL subclass. Total fatty acids also increased with higher BCS, contributing to the elevation of multiple individual fatty acid concentrations. No significant changes were observed in amino acid concentrations.

**Figure 1.**
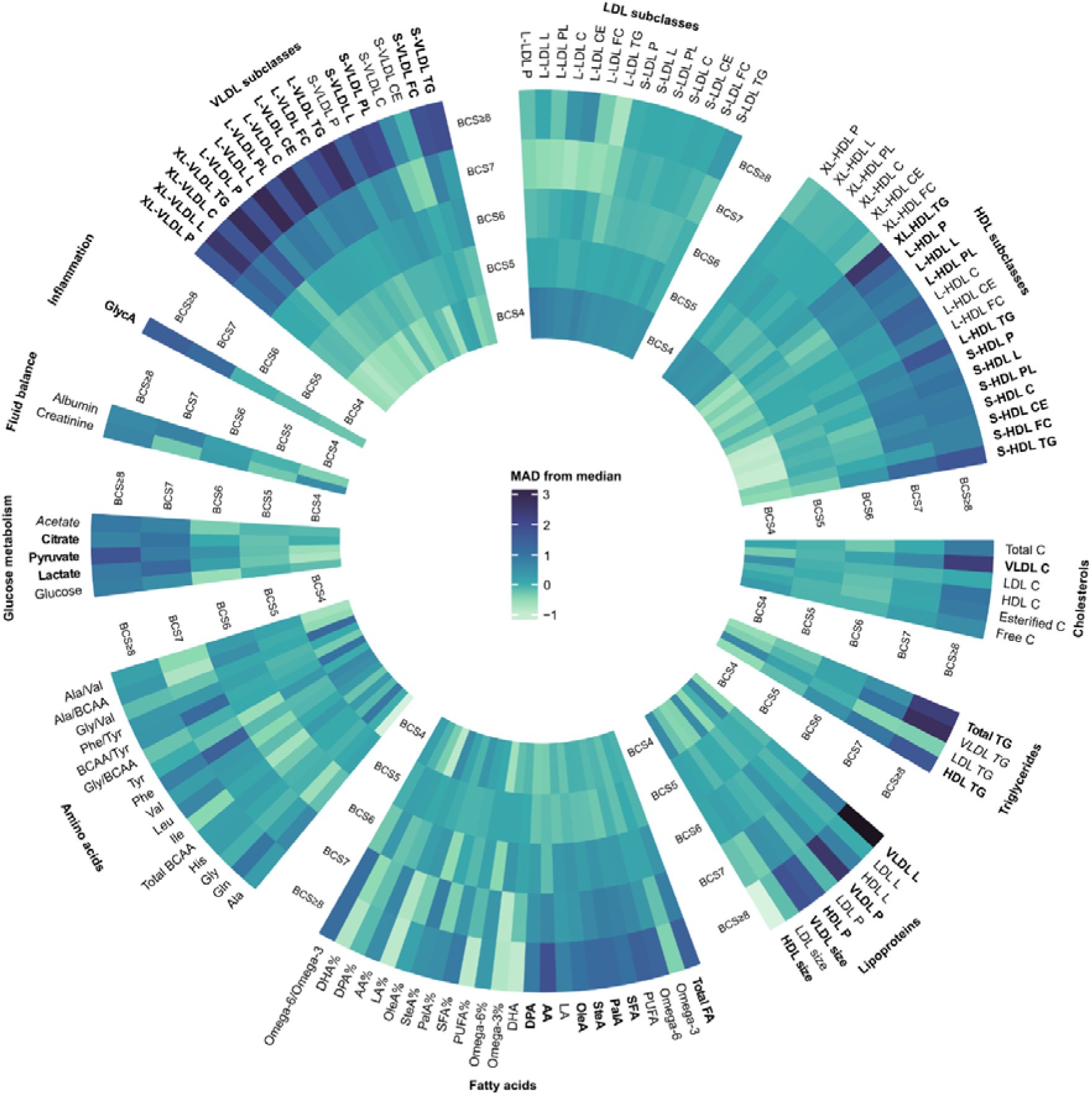
Heatmap of metabolite concentrations per BCS group. Color scaling is based on the number of median absolute deviations (MAD); the median of the BCS group differs from the median of the full dataset. Metabolites significantly (p < 0.01) associated with BCS in the generalized linear models adjusted for age and health status are emboldened. Metabolites for which reliable model creation was unsuccessful are italicized. VLDL: very low-density lipoprotein, LDL: low-density lipoprotein, HDL: high-density lipoprotein, XL-: very large, L-: large, S-: small, P: particles, L: lipids, PL: phospholipids, C: cholesterol, CE: esterified cholesterol, FC: free cholesterol, TG: triglycerides, FA: fatty acids, PUFA: polyunsaturated fatty acids, SFA: saturated fatty acids, PalA: Palmitic acid, SteA: Stearic acid, OleA: Oleic acid, LA: Linoleic acid, AA: Arachidonic acid, DPA: Docosapentaenoic acid, DHA: Docosahexaenoic acid, Ala: Alanine, Gln: Glutamine, Gly: Glycine, His: Histidine, BCAA: Branched-chain amino acids, Ile: Isoleucine, Leu: Leucine, Val: Valine, Phe: Phenylalanine, Tyr: Tyrosine, GlycA: Glycoprotein acetyls.

### Muscle condition score (MCS)

#### Demography

The demographical information of dogs included in the MCS analysis is presented in Table 2. Median age rose in all MCS groups and significant differences in age were observed between the groups. Health status was also significantly different between the MCS groups, with reversed proportions of healthy dogs and dogs having a disease in the normal muscle mass (MCS1) and moderate to severe muscle loss (MCS 3-4) groups. No other demographical characteristics were significantly different between the groups. All groups consisted of a large variety of breeds.

**Table 2.** Demographical information of the 99 dogs included in the MCS study. * significant (Kruskal-Wallis/Chi-square p < 0.05) difference between groups. ^a,b,c^: Different subscripts indicate significantly different values. MCS1: Normal muscle mass, MCS2: Mild muscle loss, MCS 3-4: moderate to severe muscle loss.

| MCS | N | Age median (range)* | %M/F | % castr | % ster | Fasting% <12h/ >12 h | % disease* | N breeds |
| --- | --- | --- | --- | --- | --- | --- | --- | --- |
| MCS1 | 44 | 9.4 (8.0 – 14.6)<sup>a</sup> | 43/57 | 42 | 64 | 41/59 | 13.6 | 30 |
| MCS2 | 40 | 11.0 (8.5 – 16.4)<sup>b</sup> | 55/45 | 36 | 78 | 52/48 | 30 | 28 |
| MCS3-4 | 15 | 12.7 (9.7 – 16.3)<sup>c</sup> | 60/40 | 44 | 83 | 53/47 | 86.7 | 12 |

No significant (chi-squared test p < 0.05) differences were observed in owner-reported daily exercise reported in hours of outdoor activity, daily exercise type, food type, how the daily food ratio was determined, how often the dog was fed treats, or how often the dog got oil supplementation. A significant difference was observed in the number of feeding times per day (once a day, twice a day, three times a day or more, ad libitum feeding, other), where dogs with moderate to severe muscle loss were more commonly fed three times or more per day.

### Effects of muscle condition on metabolism

Although the moderate to severe muscle loss group appeared visually distinct in multiple metabolite concentrations (Figure 2), many metabolite differences did not reach statistical significance (p < 0.01). Metabolic associations observed with muscle condition included increased GlycA levels, a higher phenylalanine-to-tyrosine ratio, increased concentrations of small VLDL particles, elevated lipid and cholesterol content, and heightened esterified cholesterol in large VLDL particles (Figure 2, Supplementary Table). The limited number of significant associations with MCS is primarily due to the strong interplay between muscle condition, age, and health status—particularly health status, which notably influenced various metabolites.

**Figure 2.**
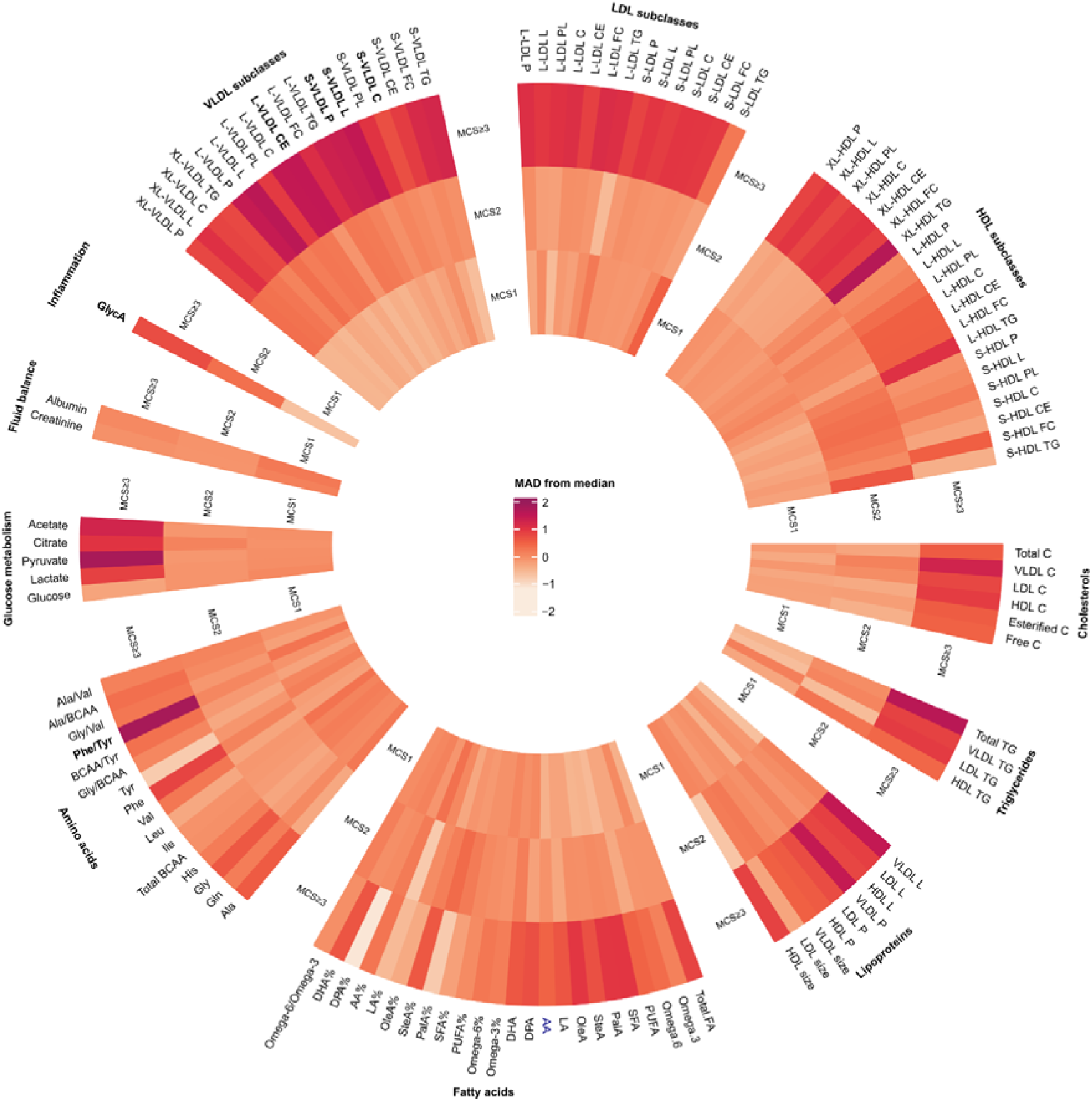
Heatmap of metabolite concentrations per MCS group in dogs over 8 years of age. Color scaling is based on the number of median absolute deviations (MAD) the median of the MCS group differs from the median of the dataset. Metabolites significantly (p < 0.01) associated with MCS in the generalized linear models are emboldened if the MCS-based model had a lower AIC than the model based on age or health status. MCS1: Normal muscle mass, MCS2: Mild muscle loss, MCS≥3: moderate to severe muscle loss. VLDL: very low-density lipoprotein, LDL: low-density lipoprotein, HDL: high-density lipoprotein, XL-: very large, L-: large, S-: small, P: particles, L: lipids, PL: phospholipids, C: cholesterol, CE: esterified cholesterol, FC: free cholesterol, TG: triglycerides, FA: fatty acids, PUFA: polyunsaturated fatty acids, SFA: saturated fatty acids, PalA: Palmitic acid, SteA: Stearic acid, OleA: Oleic acid, LA: Linoleic acid, AA: Arachidonic acid, DPA: Docosapentaenoic acid, DHA: Docosahexaenoic acid, Ala: Alanine, Gln: Glutamine, Gly: Glycine, His: Histidine, BCAA: Branched-chain amino acids, Ile: Isoleucine, Leu: Leucine, Val: Valine, Phe: Phenylalanine, Tyr: Tyrosine, GlycA: Glycoprotein acetyls.

Figure 3 illustrates p-values and the lowest AIC values from generalized linear models assessing MCS, health status, and age. Despite efforts to minimize the influence of health status on metabolite concentrations during data collection, health status remained the predominant predictor for most metabolites. In contrast, the independent effect of age on metabolite concentrations was relatively minor.

**Figure 3.**
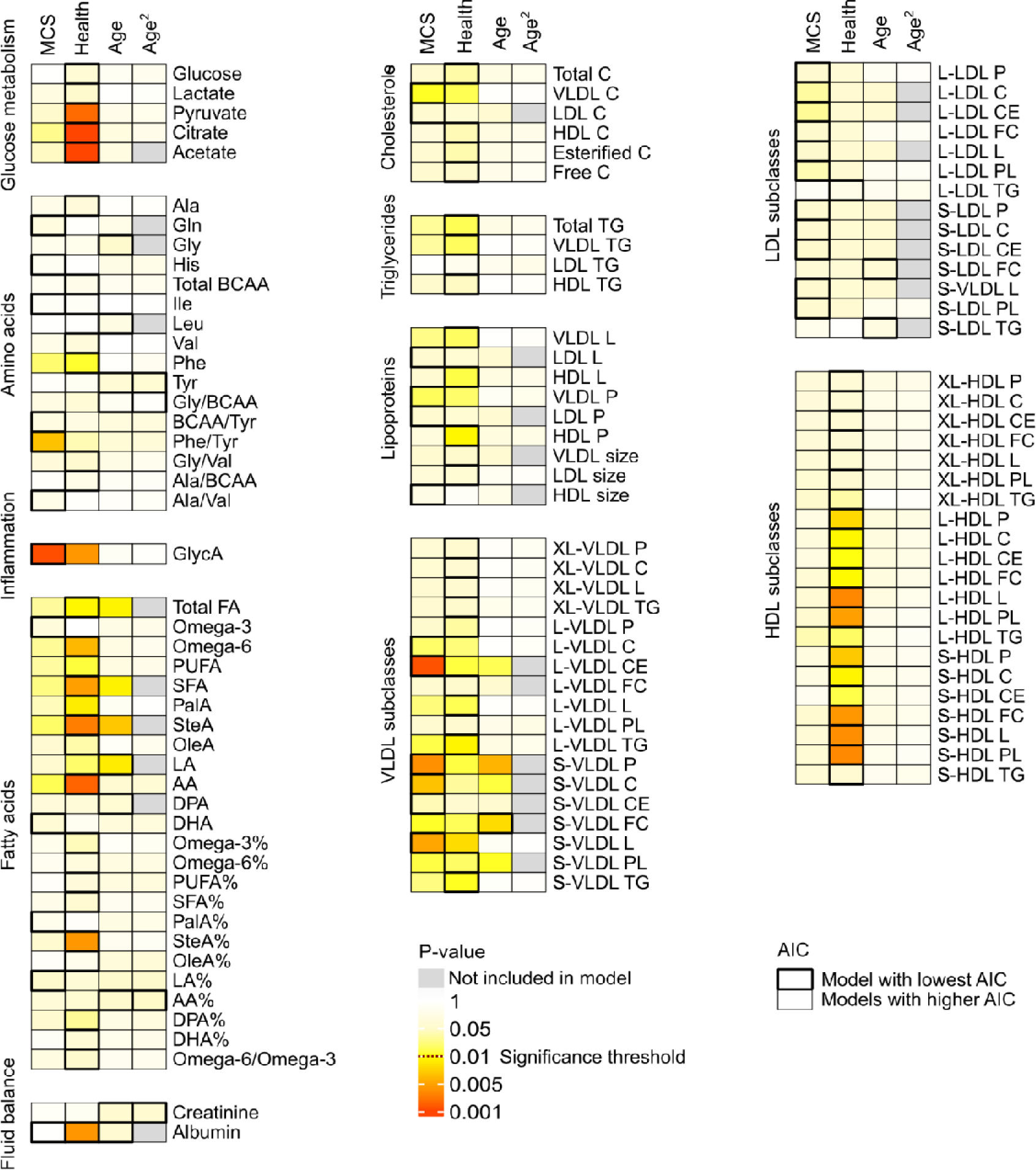
Heatmap indicating the strongness of the associations of MCS, health status, and age in dogs over 8 years of age in generalized linear models. The p-values were based on the individual generalized linear models for MCS, health status and age, except for albumin, where the full model, including all of these variables, had the lowest Akaike Information Criterion (AIC) value. For albumin, the p-values are based on the full model. Models with the lowest AIC values were considered to explain the observed changes in the metabolites’ values. Ala: Alanine, Gln: Glutamine, Gly: Glycine, His: Histidine, BCAA: Branched-chain amino acids, Ile: Isoleucine, Leu: Leucine, Val: Valine, Phe: Phenylalanine, Tyr: Tyrosine, GlycA: Glycoprotein acetyls, FA: fatty acids, PUFA: polyunsaturated fatty acids, SFA: saturated fatty acids, PalA: Palmitic acid, SteA: Stearic acid, OleA: Oleic acid, LA: Linoleic acid, AA: Arachidonic acid, DPA: Docosapentaenoic acid, DHA: Docosahexaenoic acid, C: cholesterol, TG: triglycerides, L: lipids, P: particles, VLDL: very low-density lipoprotein, LDL: low-density lipoprotein, HDL: high-density lipoprotein, XL-: very large, L-: large, S-: small, PL: phospholipids, CE: esterified cholesterol, FC: free cholesterol.

## DISCUSSION

This study identified metabolic changes associated with two significant conditions impacting canine wellbeing: i) overweight and ii) sarcopenia. This represents the largest metabolomics analysis addressing overweight conditions in dogs. The observed metabolic alterations highlight the negative effects of canine overweight, impacting various metabolite groups ranging from lipid and energy metabolism to inflammatory markers. The elevated inflammatory response noted in overweight dogs underscores the harmful effects of excessive weight on overall health, suggesting that weight reduction and metabolic monitoring could substantially enhance the animals’ wellbeing. Additionally, this study is the first to detail metabolic changes linked to reduced muscle mass in dogs. The notable associations among metabolic disturbances, muscle condition, and overall health status indicate that moderate to severe muscle loss has significant pathological implications. Early identification and monitoring of this pathological progression may substantially improve health and quality of life in aging dogs.

Associations between BCS and metabolite concentrations were linear, affecting lipid metabolism, glucose-related metabolites, and the inflammatory biomarker GlycA. These findings illustrate a continuum of dyslipidemia, elevated inflammation, and abnormal energy metabolism with increasing adiposity. Elevated plasma lactate, pyruvate, and citrate levels observed in overweight/obese dogs indicate progressively impaired glucose metabolism associated with increasing body fat. Prior studies in obese humans and dogs similarly reported elevated plasma lactate levels ^56–58^, with reduced insulin sensitivity identified as a contributing factor in humans ^58^.

The observed increase in dyslipidemia among overweight and obese dogs was anticipated, given that dyslipidemia is a well-established consequence of obesity in both dogs and humans. Numerous studies have evaluated lipid parameters in overweight or obese dogs, consistently identifying dyslipidemia despite variations in specific lipid and lipoprotein classes across studies ^56,59–65^. Differences among previous studies could be attributed to variations in lipoprotein subclass behavior and methodological approaches. Our study employed the most comprehensive lipoprotein analysis available to date, revealing specific lipid profile alterations in overweight and obese dogs, notably increased concentrations of triglycerides, fatty acids, and VLDL, alongside increased VLDL particle size, reduced HDL particle size, and elevated levels of both small and large HDL particles. Interestingly, LDL levels remained unaffected. These lipid profile changes observed in obesity differ somewhat from those associated with various diseases. For instance, inflammatory diseases typically result in increased LDL concentrations ^66^ and enlarged HDL particle size, with reduced levels of small and large HDL particles ^66^—changes opposite to those noted in obesity within this study. The clinical significance of these differing dyslipidemia profiles remains to be further elucidated.

Consistent with previous findings in both humans and dogs ^63,64,67–69^, increased overweight independently heightened the inflammatory burden, as evidenced by elevated GlycA levels. Given that obese dogs exhibit greater morbidity and mortality risks, chronic inflammation and associated metabolic disturbances likely contribute significantly to this increased risk. In humans, chronic low-grade inflammation and a compromised metabolic profile have been shown to elevate overall mortality and morbidity risk for over a decade prior to disease onset ^70–77^. This insight shifts our understanding from disease-specific etiologies toward recognizing general metabolic susceptibility, manifesting in various diseases. Similarly, in humans, obesity significantly elevates inflammatory load, and weight loss effectively reduces this inflammation ^67,68^. Consequently, weight reduction and inflammation management are crucial for improving health and longevity in overweight individuals. Monitoring inflammatory and metabolic status in overweight dogs may offer owners valuable insights into how lifestyle modifications can enhance their pet’s overall health and wellbeing.

Additionally, in this study, overweight and obese dogs more frequently suffered from underlying diseases compared to normal-weight dogs, with musculoskeletal diseases and periodontitis being the most common. This finding is particularly significant as obesity is a known risk factor for musculoskeletal disorders, often exacerbating clinical symptoms. Conversely, managing body weight may become increasingly challenging for dogs whose physical activity is limited due to existing musculoskeletal conditions.

There was a substantial difference in health status between MCS groups. While only approximately 14% of dogs in the normal muscle mass group had an underlying disease, this proportion was reversed in the moderate to severe muscle loss group. Muscle condition was closely associated with both age and health status, with health status emerging as the strongest determinant of metabolite concentrations. The high prevalence of underlying diseases and significantly altered metabolic profiles in dogs with moderate to severe muscle loss indicate that age-related sarcopenia typically results in only mild muscle loss; thus, moderate or severe muscle loss should be considered indicative of underlying disease or compromised overall health. Additionally, dogs experiencing declining MCS were fed more frequently each day compared to dogs with normal muscle mass, suggesting that owners of these dogs might be aware of their pet’s health issues and may be actively attempting to manage muscle condition, appetite, or gastrointestinal function.

The finding that health status was the primary predictor of metabolite concentrations in older dogs is particularly notable, given that this study intentionally included diseases hypothesized to have minimal metabolic impact, such as osteoarthritis, spondylosis, and periodontitis. Previously, we observed substantial metabolic alterations associated with increasing age ^25,28^, accompanied by widening confidence intervals that raised questions about whether these changes were purely age-related or attributable to subclinical diseases ^28^. The current study’s results strongly suggest that significantly abnormal metabolite values in senior dogs are more likely due to underlying disease rather than aging alone.

Independent associations between declining muscle condition and metabolic changes included elevated inflammatory burden, increased phenylalanine-to-tyrosine ratio, and mild dyslipidemia characterized by elevated cholesterol concentrations within small and large VLDL particles. In humans, sarcopenia and frailty are similarly associated with chronic low-grade inflammation ^78,79^; the observed increase in GlycA in dogs with declining muscle condition confirms this relationship in dogs as well. The elevated inflammatory burden, particularly notable in moderate to severe muscle loss, supports the pathological nature of significant muscle wasting in dogs. This chronic inflammation may contribute to increased mortality risk, as consistently demonstrated in human studies linking chronic low-grade inflammation to higher morbidity and mortality ^70–77^. Therefore, GlycA might serve as a potential biomarker for pathological muscle loss.

The increased phenylalanine-to-tyrosine ratio in dogs with poor muscle condition reflects heightened skeletal muscle catabolism, as muscle breakdown leads to an increased phenylalanine influx into the bloodstream. The phenylalanine-to-tyrosine ratio is a reliable indicator of catabolic activity ^80,81^ and has previously been associated with various inflammatory conditions in dogs ^66^. Additionally, the observed elevation in certain VLDL parameters aligns with earlier human studies that reported increased triglyceride and VLDL concentrations in sarcopenic individuals ^82,83^. Although the exact cause remains uncertain, it is hypothesized that positive energy balance—resulting from reduced physical activity and decreased basal metabolic rate—may contribute to dyslipidemia and fat accumulation in sarcopenic patients ^83^. Furthermore, skeletal muscle may independently regulate lipid metabolism ^84^, with this regulatory function diminished during sarcopenia.

Like all studies, this investigation has several limitations. Hematology and clinical chemistry parameters were unavailable for 8% of the included dogs. However, these dogs had normal physical examinations and no clinical signs of disease, making significant subclinical impacts unlikely. Exercise data relied on owner-reported durations, which may lack precision; more accurate methods such as mileage tracking or activity monitors could offer greater clarity, though detailed exercise assessment was not a primary aim of this study. While physiological variables that significantly differed between groups were statistically controlled, breed remained an uncontrolled factor. Given the large number of existing dog breeds and varied genetics in mixed breeds, completely controlling for breed would have substantially limited the study size. Sighthounds, due to their distinct metabolism, were excluded from this study ^28^.

Although the aim was to recruit at least 20 dogs per BCS and MCS group, some groups had fewer participants. Nevertheless, the statistical power remained adequate to detect significant associations between BCS, MCS, and metabolite concentrations. Additionally, random forest imputation was employed to handle missing NMR data, achieving excellent imputation performance (OOB error of 0.001). Although increasingly used in untargeted metabolomics, its suitability in targeted NMR metabolomics, as used here, remains uncertain. Future research should compare this approach to standard normalization methods to confirm it does not compromise biological or clinical interpretation.

In summary, several metabolic changes associated with BCS and MCS were identified. Increasing overweight correlated with progressive dyslipidemia, altered energy metabolism, and heightened inflammatory burden. Muscle condition was strongly associated with age and health status, with health status being the most influential determinant of metabolite levels. Independently, declining MCS was linked to increased inflammation, elevated phenylalanine-to-tyrosine ratios, and altered VLDL metabolism. These findings reinforce existing evidence regarding the negative impacts of overweight and reduced muscle mass on canine health, highlighting how metabolic status mirrors overall health and emphasizing the potential benefits of monitoring metabolic health for maintaining optimal canine wellbeing.

## ACKNOWLEDGEMENTS

Dr. Jenni Puurunen, PetMeta Labs Ltd., is thanked for contributing to the study design. Laura Parikka, Evidensia Eläinlääkäripalvelut Ltd. is thanked for the collection of study data and samples. Heta Forsström, Nightingale Health Ltd. is thanked for providing the practical needs of the study. PetMeta Labs Ltd. is thanked for enabling and funding this study. Evidensia Eläinlääkäripalvelut Ltd. is thanked for allowing sample collection in their veterinary hospitals and for scientific collaboration. The owners of the recruited dogs are thanked for the possibility of utilizing their pet dogs to advance research and dog health. The veterinarians of Eläinsairaala Evidensia Tammisto and Espoo Veterinary Hospital are thanked for their help in patient recruitment and diagnostics. Doc. Minna Rinkinen at Evidensia Eläinlääkäripalvelut Ltd. is thanked for checking the written MCS class descriptions utilized in the MCS evaluation.

## Author contribution

Claudia Ottka: Conceptualization and study design. Conduction of statistical analyses. Interpreting the results and drafting the manuscript.

Mirja Kaimio: Supervising and practical planning of the study in Evidensia premises. Commenting and approving the manuscript.

Essi Leminen: Inclusion of study participants, collecting samples and data for the study. Commenting and approving the manuscript.

Hannes Lohi: Conceptualization, design and supervision of the study. Editing the manuscript.

